# A 63-bp insertion in exon 2 of the porcine *KIF21A* gene is associated with arthrogryposis multiplex congenita

**DOI:** 10.1101/2020.04.09.033761

**Authors:** Zih-Hua Fang, Adéla Nosková, Danang Crysnanto, Stefan Neuenschwander, Peter Vögeli, Hubert Pausch

## Abstract

Arthrogryposis multiplex congenita (AMC) is a recessively inherited fatal disease detected almost 20 years ago in the Swiss Large White pig population. A diagnostic marker test enabled the identification of carrier animals, but the underlying causal mutation remains unknown. To identify the mutation underlying AMC, we collected whole-genome genotyping and sequencing data for 11 affected piglets and 23 healthy pigs. Haplotype-based case-control association testing using 47,829 SNPs confirmed that AMC maps to SSC5 (*P* = 9.4 ×10^−13^). Subsequent autozygosity mapping revealed a common 6.06 Mb region (from 66,757,970 to 72,815,151 bp) of extended homozygosity in 11 piglets affected by AMC. We detected a 63-bp insertion in the second exon of *KIF21A* gene encoding Kinesin Family Member 21A using whole-genome sequences of a carrier boar, two of its affected and two heterozygous piglets. This insertion was compatible with the recessive inheritance of AMC. The 63-bp insertion likely represents a loss-of-function allele because it is predicted to introduce a premature stop codon in *KIF21A* gene (p.Val41_Phe42insTer) that truncates 1,614 amino acids (∼ 97%) from the protein. Lack of KIF21A protein is lethal in mice, thus providing additional evidence that a loss-of function allele of *KIF21A* might cause fatal AMC in pigs. We found that this deleterious allele still segregates at low frequency in the Swiss Large White pig population. The unambiguous detection of carrier animals can now facilitate the eradication of the deleterious allele from the population.

---

Small effective population size and the wide-spread use of a few breeding animals may result in the frequent phenotypic manifestation of recessive alleles in livestock populations. Microarray-based genotyping in affected and unaffected individuals facilitates rapid mapping of recessive genetic defects (Charlier *et al*. 2008). The analysis of whole-genome sequence variant genotypes of either affected or carrier animals has proven to be very efficient in detecting causal variants for recessive traits (https://omia.org/home/). Yet uncovering causal variants remains challenging when they reside in genomic regions with either assembly problems or gaps in reference sequences (Pausch *et al*. 2015). Diagnostic haplotype- or microsatellite-based testing may be used to detect carrier animals and avoid at-risk matings even when the molecular basis of genetic defects is unknown. However, the presence of either recombinant or ancestral haplotypes may prevent unambiguous identification of carrier animals when causal mutations are not genotyped (e.g. Jung *et al*. 2014).

Arthrogryposis multiplex congenita (AMC) is a fatal autosomal recessive disease that was first described almost 20 years ago in the Swiss Large White pig population (Genini *et al*. 2004). Affected piglets die either shortly before or during birth and show various congenital abnormalities including malformed and contracted joints of fore- and hindlimbs, brachygnathia inferior and scoliosis (Genini *et al*. 2004, Haubitz *et al*. 2012). Using linkage analysis, a 2.32 Mb region (1.86 Mb based on the Sscrofa 11.1 assembly) of SSC5 has been associated with AMC (Haubitz *et al*. 2012). Microsatellite testing was applied to exclude carrier animals from breeding and remove the deleterious allele from the population. However, a causal mutation had not been detected so far. Here, we attempt to eventually identify the causal mutation for AMC using whole-genome genotyping and sequencing data.

Initially, Genini *et al*., 2004 mapped AMC to a 4.61 Mb interval on SSC5 between two microsatellite markers SW152 (69,619,907 bp, Sscrofa 11.1) and SW904 (74,230,251 bp, Sscrofa 11.1) using linkage analysis. Using a larger dataset and additional markers enabled refining the AMC-associated region to a 1.86 Mb interval between two SNP markers ALGA0032767 (70,454,750 bp, Sscrofa 11.1) and DRGA0006010 (72,315,123 bp, Sscrofa 11.1; Haubitz *et al*. 2012). To confirm the genomic region associated with AMC, we genotyped cases and controls using Illumina PorcineSNP60 Genotyping Bead chips (see data availability). To do so, we isolated genomic DNA of 11 affected piglets and 23 healthy pigs for genotyping at 62,543 SNPs using either frozen EDTA-stabilized blood or frozen tail biopsies. The physical positions of the SNPs were based on the Sscrofa11.1 assembly of the porcine genome (Warr *et al*. 2019). Quality control was carried out using PLINK (version 1.9; Chang *et al*. 2015). We retained 47,829 autosomal SNPs that were genotyped in more than 90% of the samples, had MAF greater than 0.01 and did not deviate significantly from Hardy-Weinberg proportions (*P*-value greater than 1 ×10^− 4^). To impute sporadically missing genotypes and infer haplotypes, we applied 25 iterations of the phasing algorithm implemented in Beagle 5.0 (Browning *et al*. 2018) assuming an effective population size of 200 and all other parameters set to default values. Subsequently, we applied a sliding window-based approach to compare haplotype frequency in cases and controls as described in Venhoranta *et al*. 2014 and Pausch *et al*. 2016. Specifically, we shifted a sliding window consisting of 20 contiguous SNPs along the chromosomes in steps of 5 SNPs. Within each window, haplotypes with frequency greater than 1% were tested for association with AMC using Fisher’s exact test of allelic association.

The strongest association signal resulted from six adjacent haplotype windows with identical *P* values (*P* = 9.4 ×10^−13^) that were located on SSC5 between 69,601,646 and 71,330,075 bp (Fig.1a). The top haplotypes partly overlap the AMC-associated interval defined by Haubitz *et al*. (2012). The top haplotypes are in the center of a 6.06 Mb segment (from 66,757,970 to 72,815,151 bp) of extended homozygosity that was identical by descent in the 11 piglets affected by AMC. This segment was not detected in the homozygous state in 23 healthy pigs (Fig. 1b). According to the Ensembl (version 98) annotation of the porcine genome, the segment encompasses 67 genes. No haplotypes located outside the 6.06 Mb segment of extended homozygosity were associated with AMC at a Bonferroni-corrected significance threshold of 4.0 ×10^−7^.

**Fig. 1.**
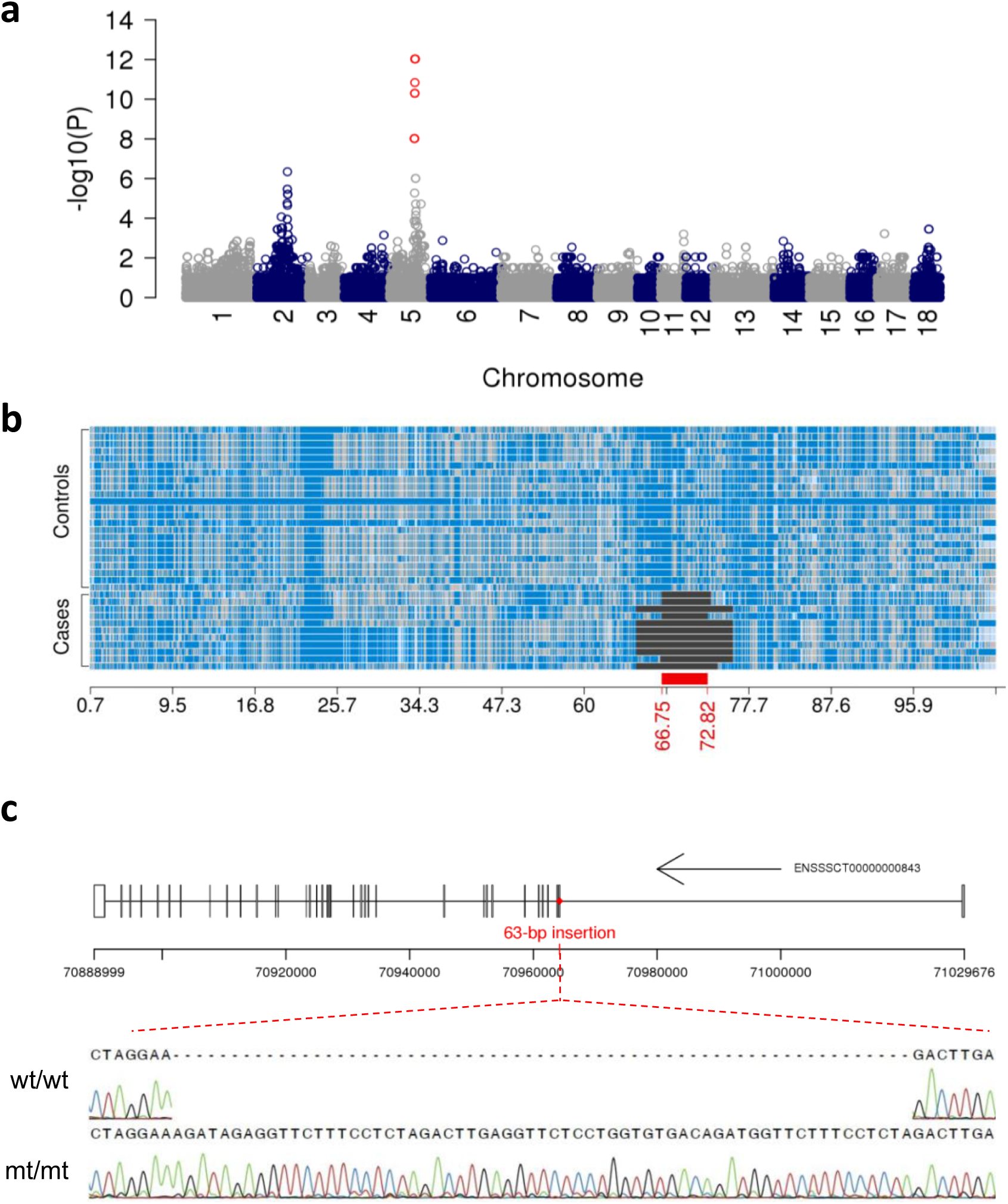
Identification of a 63-bp insertion in exon 2 of porcine *KIF21A* gene. (a). Manhattan plot representing the result of a haplotype-based genome-wide association study. Red color represents haplotypes with a *P* value less than 4.03 ×10^−7^ (Bonferroni corrected-significance threshold). (b). Homozygosity mapping in 11 cases and 23 controls using SNPs located on SSC5. Blue and pale blue represent homozygous genotypes (AA and BB), heterozygous genotypes are displayed in light grey. The red bar indicates a 6.06 Mb segment of extended homozygosity shared in 11 cases. (c). The 63-bp insertion is located in exon 2 (ENSSSCE00000006489) of *KIF21A* (transcript ID: ENSSSCT00000000843). Sanger sequencing chromatograms from one wild type (wt/wt) pig and one mutant (mt/mt) piglet.

Next, we sequenced a carrier boar, two of its affected and two heterozygous piglets to an average of 13.2-fold read depth using 2 ×150 bp paired-end reads (see data availability). Illumina TruSeq PCR-free libraries with insert sizes of 350 bp were prepared and sequenced with an Illumina NovaSeq6000 instrument. We used the fastp software (Chen *et al*. 2018) to remove adapter sequences and reads that had phred-scaled quality less than 15 for more than 15% of the bases. Subsequently, the filtered reads were aligned to the Sscrofa 11.1 assembly of the porcine genome using the BWA-MEM algorithm implemented in the BWA software (Li 2013). The Picard tools software suite (Picard Toolkit, 2019) and Sambamba (Tarasov *et al*. 2015) were applied to mark duplicates and sort the alignments by coordinates, respectively. The variant detection (SNPs and Indels) in the five sequenced animals was performed together with 93 pigs for which sequence data were also available from our in-house database using the multi-sample variant calling approach implemented in the Genome Analysis Toolkit (GATK, version 4.1.0; Depristo *et al*. 2011). Subsequently, we filtered the detected sequence variants by hard-filtering according to GATK’s best practice guidelines. The retained sequence variants were annotated using the Ensembl Variant Effect Predictor (VEP, release 98; McLaren *et al*. 2016). We also discovered and genotyped large structural variants (> 50 bp) including insertions, deletions, inversions, duplications and translocations using Pindel v0.2.5b9 (Ye *et al*. 2009) and Delly v0.7.8 (Rausch *et al*. 2012) with the default settings. However, we did not detect structural variants (> 50 bp) that were compatible with recessive inheritance of AMC. The analysis of sequencing depth along chromosome 5 in affected and unaffected pigs using mosdepth (Pedersen and Quinlan 2018) did not reveal large deletions or duplications segregating with the AMC-associated haplotype.

To detect candidate causal variants for AMC, we considered 75,110 sequence variants located within the 6.06 Mb segment of extended homozygosity on SSC5 (from 66,757,970 to 72,815,151 bp). We retained variants as candidate causal mutations if they were homozygous in two affected piglets, heterozygous in the carrier boar and two carrier piglets, and not homozygous in 93 AMC-unaffected pigs for which the haplotype status was unknown. This filtering identified 809 variants that were compatible with recessive inheritance (see data availability). Of these 809 variants, only two were predicted to alter protein-coding sequences: a 6-bp in-frame insertion (NC_010447.5:g71673414_71673414insCGCCCG, p.Ala56_Gly57dup) in *SLC2A13* gene encoding Solute Carrier Family 2 Member 13 that was predicted to have a moderate impact on the resulting protein, and a 63-bp insertion in *KIF21A* gene encoding Kinesin Family Member 21A that was predicted to introduce premature termination of translation (NC_010447.5:g70964237_ g70964238insAAGATAGAGGTTCTTTCCTCTAGACTTGAGGTTCTCCTGGTGTGA CAGATGGTTCTTTCCTCT, p.Val41_Phe42insTer). Furthermore, 807 out of 809 variants were detected in the heterozygous state in at least five AMC-unaffected control pigs from our in-house database for which the haplotype status was unknown, indicating that they occur at least at low to moderate frequency in the population. Only two variants were never detected either in the heterozygous or homozygous state in control pigs: the 63-bp insertion in *KIF21A* gene and a variant (NC_010447.5:g71800313) in the 5’UTR of *LRKK2*, further suggesting that the 63-bp insertion is a plausible causal mutation for AMC. We confirmed the presence of the 63-bp insertion in *KIF21A* gene in 11 affected piglets using PCR and subsequent sanger sequencing of PCR products from one affected piglet and one homozygous wildtype pig (Table S1, Fig.1c).

To investigate if the AMC-associated allele still segregates unnoticed in the Swiss Large White population, we screened genotypes (Porcine 60K SNP Bead chips) of 9,206 healthy pigs available from routine genomic breeding value estimation for the presence of the AMC-associated haplotype. This haplotype was in perfect LD with the 63-bp insertion in our AMC pedigree of 11 affected piglets and 23 unaffected pigs. The frequency of the AMC-associated haplotype was 1.61% in the Swiss Large White pig population. We detected the AMC-associated haplotype in 293 pigs in the heterozygous state and, surprisingly, in two healthy pigs in the homozygous state. We obtained biological materials (blood, ear tissue or hair samples) of 30 randomly selected heterozygous haplotype carriers and one homozygous haplotype carrier to isolate genomic DNA for testing the presence of the 63-bp insertion. Biological material was not available for the second homozygous haplotype carrier. PCR testing revealed that only 2 (out of 30) heterozygous haplotype carriers also carried the 63-bp insertion, indicating that either an ancestral or recombinant haplotype without the insertion also segregates in the population. The one homozygous haplotype carrier did not carry the 63-bp insertion in *KIF21A* gene. Assuming that 7% (2/30) of the haplotype carriers are indeed mutation carriers, the 63-bp insertion has an allele frequency of approximately 0.1% in the Swiss Large White population.

The *KIF21A* gene encodes Kinesin Family Member 21A, a microtube-dependent motor protein that is predominantly expressed in the central nervous system and muscle (Marszalek *et al*. 1999, Desai *et al*. 2012). Kinesin Family Member 21A plays a crucial role in cell morphogenesis and neuronal development by regulating microtubule growth (van der Vaart *et al*. 2013). The premature stop codon in porcine *KIF21A* truncates 1,614 amino acids (∼ 97%) from KIF21A (transcript ID: ENSSSCT00000000843). If not degraded via nonsense-mediated mRNA decay, the truncated protein lacks a complete motor domain (Yamada *et al*. 2003). Mice lacking the motor domain of KIF21A die within 24h of birth (Cheng *et al*. 2014). Piglets homozygous for the 63-bp insertion were either stillborn or died shortly after birth and had congenital malformations, suggesting that a lack of the KIF21A motor domain is also lethal in the homozygous state in pigs. The multi-species comparison of KIF21A protein sequences using the constraint-based alignment tool (COBALT; Papadopoulos and Agarwala, 2007) shows a strong evolutionarily conservation of the motor domain (position 9-371), further suggesting its importance in the physiological protein function (Fig. S1).

Heterozygous missense mutations in human *KIF21A* gene impair the function of the oculomotor nerve due to fibrosis of extraocular muscles (Yamada *et al*. 2003, Lu *et al*. 2008, Bianchi *et al*. 2016). Apart from eye movement disorders, people with pathogenic *KIF21A* mutation are healthy. Pigs carrying the AMC-associated haplotype harboring the 63-bp insertion in the heterozygous state were also healthy. However, their eye movement had not been investigated and described in detail. Fibrotic processes may possibly underlie the characteristically contracted joints observed in the piglets with AMC (Kalampokas *et al*. 2012). Interestingly, loss-of-function mutations of human *KIF21A* have not been observed in the homozygous state so far, suggesting that they might be lethal (Walsh and Engle 2010). We did not detect the 63-bp insertion in *KIF21A* in the homozygous state, suggesting that it represents a lethal allele. Furthermore, mutations in kinesin superfamily proteins (KIFs) cause spasticity of limbs (Reid *et al*. 2002, Dor *et al*. 2014, Duchesne *et al*. 2018), supporting our findings that a loss-of-function allele in porcine *KIF21A* gene is associated with AMC.

The identification of a plausible causal mutation for a recessive disease emerged almost 20 years ago evidences that current whole-genome genotyping and sequencing technologies may readily reveal plausible candidate causal mutations for unresolved genetic disorders. However, our results do not provide functional evidence that the putative loss-of-function allele in *KIF21A* is causal for AMC. Porcine AMC was investigated decades ago, and tissue samples of affected piglets had not been preserved once diagnostic marker testing indicated that the deleterious allele was eradicated from the population. Due to the lack of appropriate tissue samples, we could not investigate either mRNA or protein expression in affected piglets to validate the consequence of the 63-bp insertion in *KIF21A* gene. Nevertheless, our findings indicate that deleterious alleles of *KIF21A* warrant close scrutiny in unresolved cases of neurological and muscular disorders.

In conclusion, we report a plausible candidate causal variant for AMC in Swiss Large White pigs. A 63-bp insertion in *KIF21A* gene likely represents a loss-of-function allele that causes this fatal disease in the homozygous state.

## Supporting information

Table S1 PCR primers and condition for genotyping and sanger sequencing the 63-bp insertion in porcine KIF21A gene

Fig. S1 Alignment of KIF21A protein sequences in multiple species using the constraint-based alignment tool (COBALT).

## Acknowledgements

We acknowledge SUISAG for providing genotype data and biological materials of healthy control animals. The sequencing of unaffected control animals was financially supported by SUISAG, Micarna SA and the ETHZ Foundation.

## Availability of data

Whole-genome sequences are available at the Sequence Read Archive of the NCBI at the BioProject PRJNA622908 under sample accession numbers: SAMN14532191 and SAMN14532478 (2 piglets affected with AMC), SAMN14532792 (the carrier boar), and SAMN14532769 and SAMN14532790 (two heterozygous piglets). Whole-genome genotyping data of 11 affected and 23 healthy animals together with a VCF file containing 809 sequence variants compatible with recessive inheritance are available at https://doi.org/10.5281/zenodo.3746178.

## Supporting information

Table S1 PCR primers and condition for genotyping and sanger sequencing the 63-bp insertion in porcine *KIF21A* gene

## Notes

### Competing Interest Statement

The authors have declared no competing interest.

