## Supplementary material for "A 63-bp insertion in exon 2 of the porcine *KIF21A* gene is associated with arthrogryposis multiplex congenita": Fig. S1 Alignment of KIF21A protein sequences in multiple species using the constraint-based alignment tool (COBALT).

```
NP_001166935.1 (Home sapiens)        MLGAPDE--SSVRVAV-------RIRPQLAKEKIEGCHICTSVTPGEPQVFLGKDKAFTFDYVFDIDSQQEQIYIQCIEK  71
XP_017171930.1 (Mus musculus)        -MGPATWacVHGCWRVrsydpwsRIRPQLAKEKIEGCHICTSVTPGEPQVFLGKDKAFTFDYVFDIDSQQEQIYTQCIEK  79
XP_017450778.1 (Rattus norvegicus)   MLGAADEssV----RVav-----RIRPQLAKEKIEGCHICTSVTPGEPQVFLGKDKAFTFDYVFDIDSQQEQIYTQCIEK  71
NP_001193847.1 (Bos Taurus)          MWGAPDE--SSVRVAV-------RIRPQLAKERIEGCHICTSVTPGEPQVFLGKDKAFTFDYVFDIDSQQEQIYIQCIEK  71
NP_001182593.1 (Xenopus tropicalis)  MTGAQDD--SAVRVAL-------RIRPQLAKEKIEGCHICTSVTPGEPQVVLGKDKAFTFDYVFDIDSRQEEIYMQCTEK  71
XP_016779133.1 (Pan troglodytes)     MLGAPDE--SSVRVAV-------RIRPQLAKEKIEGCHICTSVTPGEPQVFLGKDKAFTFDYVFDIDSQQEQIYIQCIEK  71
XP_005637036.2 (Canis lupus)         MFFITQH--VTSCMWIittellsRIRPQLAKEKIEGCHICTSVTPGEPQVFLGKDKAFTFDYVFDIDSQQEQIYTQCIEK  78
XP_013853381.2 (Sus Scrofa)          MWGAPDE--SSVRVAV-------RIRPQLAKERIEGCHICTSVTPGEPQVFLGKDKAFTFDYVFDIDSQQEQIYTQCIEK  71
XP_027823250.1 (Ovis aries)          MWGAPDE--SSVRVAV-------RIRPQLAKERIEGCHICTSVTPGEPQVFLGKDKAFTFDYVFDIDSQQEQIYTQCIEK  71
XP_017903411.1 (Capra hircus)        MWGAPDE--SSVRVAV-------RIRPQLAKERIEGCHICTSVTPGEPQVFLGKDKAFTFDYVFDIDSQQEQIYTQCIEK  71
XP_015131286.1 (Gallus gallus)       MSAGQDE--SSVRVAV-------RIRPQLAKEKIEGCHICTSVTPGEPQVFLGKDKAFTFDYVFNIDSQQEEIYIQCIEK  71

NP_001166935.1 (Home sapiens)        LIEGCFEGYNATVFAYGQTGAGKTYTMGTGFDVNIVEEELGIISRAVKHLFKSIEEKKHIAIKNGLPAPDFKVNAQFLEL  151
XP_017171930.1 (Mus musculus)        LIEGCFEGYNATVFAYGQTGAGKTYTMGTGFDVNIMEEEQGIISRAVRHLFKSIDEKKTSAIKNGLPPPEFKVNAQFLEL  159
XP_017450778.1 (Rattus norvegicus)   LIEGCFEGYNATVFAYGQTGAGKTYTMGTGFDVNIMEEEQGIISRAVKHLFKSIEEKKTSAIKNGVPPPEFKVNAQFLEL  151
NP_001193847.1 (Bos Taurus)          LIEGCFEGYNATVFAYGQTGAGKTYTMGTGFDVNIIEEEQGIISRAVKHLFKSIEEKKHTSIKNGLPPPDFKVNAQFLEL  151
NP_001182593.1 (Xenopus tropicalis)  LIEGCFEGYNATVFAYGQTGAGKTYTMGTGFDVNISEEEHGIIPRAVTHLFRRIEEKKQFALEQGLPPPDFKVNAQFLEL  151
XP_016779133.1 (Pan troglodytes)     LIEGCFEGYNATVFAYGQTGAGKTYTMGTGFDVNIVEEELGIISRAVKHLFKSIEEKKHIAIKNGLPAPDFKVNAQFLEL  151
XP_005637036.2 (Canis lupus)         LIEGCFEGYNATVFAYGQTGAGKTYTMGTGFDVNIIEEEQGIISRAVKHLFKSIEEKKHTAIKNGHPPPDFKVNAQFLEL  158
XP_013853381.2 (Sus Scrofa)          LIEGCFEGYNATVFAYGQTGAGKTYTMGTGFDVNIIEEEQGIISRAVKHLFKSIEEKKHASIKNGLPSPDFKVNAQFLEL  151
XP_027823250.1 (Ovis aries)          LIEGCFEGYNATVFAYGQTGAGKTYTMGTGFDVNIIEEEQGIISRAVKHLFKSIEEKKHTSIKNGLPPPDFKVNAQFLEL  151
XP_017903411.1 (Capra hircus)        LIEGCFEGYNATVFAYGQTGAGKTYTMGTGFDVNIIEEEQGIISRAVKHLFKSIEEKKHTSIKNGLPPPDFKVNAQFLEL  151
XP_015131286.1 (Gallus gallus)       LIEGCFEGYNATVFAYGQTGAGKTYTMGTGFDVNITEEEQGIISRAVKHLFRCIEEKKQAAIKQGLPPPDFKVNAQFLEL  151

NP_001166935.1 (Home sapiens)        YNEEVLDLFDTTRDIDAKSKKSNIRIHEDSTGGIYTVGVTTRTVNTESEMMQCLKLGALSRTTASTQMNVQSSRSHAIFT  231
XP_017171930.1 (Mus musculus)        YNEEVLDLFDTTRDIDAKNKKSNIRIHEDSTGGIYTVGVTTRTVNTEPEMMQCLKLGALSRTTASTQMNVQSSRSHAIFT  239
XP_017450778.1 (Rattus norvegicus)   YNEEVLDLFDTTRDIDAKNKKSNIRIHEDSTGGIYTVGVTTRTVNTEPEMMQCLKLGALSRTTASTQMNVQSSRSHAIFT  231
NP_001193847.1 (Bos Taurus)          YNEEVLDLFDTTRDIDAKTKKSNIRIHEDSAGGIYTVGVTTRTVNTESEMMQCLKLGALSRTTASTQMNVQSSRSHAIFT  231
NP_001182593.1 (Xenopus tropicalis)  YNEEVLDLFDTTRDIDARNKKSNIRIHEDSSGGIYTVGVTTRNVSSETEMIQCLKIGALSRTTASTQMNVQSSRSHAIFT  231
XP_005637036.2 (Pan troglodytes)     YNEEVLDLFDTTRDIDAKSKKSNIRIHEDSTGGIYTVGVTTRTVNTESEMMQCLKLGALSRTTASTQMNVQSSRSHAIFT  231
XP_013853381.2 (Canis lupus)         YNEEVLDLFDTTRDIDAKNKKSNIRIHEDSTGGIYTVGVTTRTVNTESEMMQCLKLGALSRTTASTQMNVQSSRSHAIFT  238
XP_027823250.1 (Sus Scrofa)          YNEEVLDLFDTTRDIDAKNKKSNIRIHEDSAGGIYTVGVTTRTVNTESEMMQCLKLGALSRTTASTQMNVQSSRSHAIFT  231
XP_027823250.1 (Ovis aries)          YNEEVLDLFDTTRDIDAKTKKSNIRIHEDSAGGIYTVGVTTRTVNTESEMMQCLKLGALSRTTASTQMNVQSSRSHAIFT  231
XP_017903411.1 (Capra hircus)        YNEEVLDLFDTTRDIDAKTKKSNIRIHEDSAGGIYTVGVTTRTVNTESEMMQCLKLGALSRTTASTQMNVQSSRSHAIFT  231
XP_015131286.1 (Gallus gallus)       YNEEILDLFDTTRDIDAKNKKSNIKIHEDSAGGIYTVGVTTRTVNGESEMMQCLKLGALSRTTASTQMNVQSSRSHAIFT  231
```

```
NP_001166935.1 (Home sapiens)         IHVCQTRVCPQIDADNATDNKIISESAQMNEFETLTAKFHFVDLAGSERLKRTGATGERAKEGISINCGLLALGNVISAL  311
XP_017171930.1 (Mus musculus)         IHVCQTRVCPQTDAENATDNKLISESSPMNEFETLTAKFHFVDLAGSERLKRTGATGERAKEGISINCGLLALGNVISAL  319
XP_017450778.1 (Rattus norvegicus)    IHVCQTRVCPQTDAENVTDNKMVSESPQMNEFETLTAKFHFVDLAGSERLKRTGATGERAKEGISINCGLLALGNVISAL  311
NP_001193847.1 (Bos Taurus)           IHLCQTRMCPQIDAEIATDNKVISESSQMNEFETLTAKFHFVDLAGSERLKRTGATGERAKEGISINCGLLALGNVISAL  311
NP_001182593.1 (Xenopus tropicalis)   IHLCQNRVCPKIDNENDLDNRMASESNQINEFETLTAKFHFVDLAGSERLKRTGATGERAKEGISINCGLLALGNVISAL  311
XP_005637036.2 (Pan troglodytes)      IHVCQTRVCPQIDADNATDNKITSESAQMNEFETLTAKFHFVDLAGSERLKRTGATGERAKEGISINCGLLALGNVISAL  311
XP_013853381.2 (Canis lupus)          IHLCQTRMCPQTDAENATDNKVISESSQLNEFETLTAKFHFVDLAGSERLKRTGATGERAKEGISINCGLLALGNVISAL  318
XP_027823250.1 (Sus Scrofa)           IHLSQTRMCPQIDTENAIDNKVISESSQMNEFETLTAKFHFVDLAGSERLKRTGATGERAKEGISINCGLLALGNVISAL  311
XP_027823250.1 (Ovis aries)           IHLCQTRMCPQIDAESATDNKVISESSQMNEFETLTAKFHFVDLAGSERLKRTGATGERAKEGISINCGLLALGNVISAL  311
XP_017903411.1 (Capra hircus)         IHLCQTRMCPQIDAESATDNKVISESSQMNEFETLTAKFHFVDLAGSERLKRTGATGERAKEGISINCGLLALGNVISAL  311
XP_015131286.1 (Gallus gallus)        IHLCQTRVCPAFNTDNATDNRIISESSEMNEFETLTAKFHFVDLAGSERLKRTGATGERAKEGISINCGLLALGNVISAL  311

NP_001166935.1 (Home sapiens)         GDKSKRATHVPYRDSKLTRLLQDSLGGNSQTIMIACVSPSDRDFMETLNTLKYANRARNIKNKVMVNQDRASQQINALRS  391
XP_017171930.1 (Mus musculus)         GDKSKRATHVPYRDSKLTRLLQDSLGGNSQTIMIACVSPSDRDFMETLNTLKYANRARNIKNKVMVNQDRASQQINALRS  399
XP_017450778.1 (Rattus norvegicus)    GDKSKRATHVPYRDSKLTRLLQDSLGGNSQTIMIACVSPSDRDFMETLNTLKYANRARNIKNKVMVNQDRASQQINALRS  391
NP_001193847.1 (Bos Taurus)           GDKSKRATHVPYRDSKLTRLLQDSLGGNSQTIMIACVSPSDRDFMETLNTLKYANRARNIKNKVMVNQDRASQQINALRS  391
NP_001182593.1 (Xenopus tropicalis)   GDKSKKATHVPYRDSKLTRLLQDSLGGNSQTVMIACVSPSDRDFMETLNTLKYANRARNIKNKVMVNQDRTSQQINALRN  391
XP_005637036.2 (Pan troglodytes)      GDKSKRATHVPYRDSKLTRLLQDSLGGNSQTIMIACVSPSDRDFMETLNTLKYANRARNIKNKVMVNQDRASQQINALRS  391
XP_013853381.2 (Canis lupus)          GDKSKRATHVPYRDSKLTRLLQDSLGGNSQTIMIACVSPSDRDFMETLNTLKYANRARNIKNKVMVNQDRASQQINALRS  398
XP_027823250.1 (Sus Scrofa)           GDKSKRATHVPYRDSKLTRLLQDSLGGNSQTIMIACVSPSDRDFMETLNTLKYANRARNIKNKVMVNQDRASQQINALRN  391
XP_027823250.1 (Ovis aries)           GDKSKRATHVPYRDSKLTRLLQDSLGGNSQTIMIACVSPSDRDFMETLNTLKYANRARNIKNKVMVNQDRASQQINALRS  391
XP_017903411.1 (Capra hircus)         GDKSKRATHVPYRDSKLTRLLQDSLGGNSQTIMIACVSPSDRDFMETLNTLKYANRARNIKNKVMVNQDRASQQINALRS  391
XP_015131286.1 (Gallus gallus)        GDKSKKATHVPYRDSKLTRLLQDSLGGNSQTLMIACVSPSDRDFMETLNTLKYANRARNIKNKVMVNQDRASQQINALRN  391
```
